# Induction of osteogenic differentiation of MSCs by GSK3β knockdown through GSK3β siRNAs transfection

**DOI:** 10.1101/2023.03.21.533598

**Authors:** Elena V. Galitsyna, Anastasiia A. Buianova, Tatiana B. Bukharova, Irina A. Krivosheeva, Mikhail Yu. Skoblov, Dmitriy V. Goldshtein

**Affiliations:** Research Centre for Medical Genetics, 1 Moskvorechye Str., 115522 Moscow, Russia

**Keywords:** siRNAs, GSK3β, AD-MSCs, SHEDs, MSCs osteogenic differentiation, MSCs transfection, PEI

## Abstract

The development of effective strategies for the treatment of bone defects is based on gene therapy methods aimed at regulating the differentiation of osteoprogenitor cells. One approach is the development of knockdown systems of inhibitory genes of osteogenic cell differentiation using siRNA molecules. In this work, we developed approaches to induce osteogenic differentiation of mesenchymal stem cells (MSCs) by knockdown of *GSK3β* using siRNAs in cultures of MSCs derived from human adipose tissue (AD-MSCs). For this purpose, we performed a comparative evaluation of the efficacy of lipoplexes and polyplexes formed with one of the 4 siRNA molecules and 5 commercial transfection agents most commonly used in laboratory practice. The most effective transfection agent appeared to be PEI, which demonstrated high cytocompatibility in free form and as part of polyplexes even when maximum concentrations were used. Using the polyplexes formed by siRNA molecule designed for the first time and PEI, we developed a highly efficient *GSK3β* gene knockdown system, which showed its effectiveness in cultures of AD-MSCs. As a result, we demonstrated the osteoinductive properties of GSK3β siRNA molecules in these cultures. The results obtained can be applied in the development of gene therapy strategies based on siRNA molecules in human bone tissue diseases.

## 1. Introduction

The use of small interfering RNA (siRNA) molecules for organotypic bone tissue regeneration is a new and potentially convenient tool for regenerative medicine. Mesenchymal stem cells (MSCs) are one of the main *in vitro* models for the development of techniques in the field of gene-cell technology due to their easy derivation from different tissues and high proliferative and differentiation potential [1].

Positive regulation of osteogenic differentiation of mammalian stem cells can be achieved by both known osteoinducers – recombinant proteins of Bone morphogenetic protein (BMP) and Wingless/Integrated protein (WNT) families, dexamethasone, sodium β-glycerophosphate, calcium poly(ethylene phosphate)s, L-ascorbic acid and metabolites of vitamin D [2–6], and as a result of suppression of protein activity or expression of osteogenesis inhibitor genes by small molecules or siRNA molecules [7–9]. However, the regulation of signaling pathways by the above-mentioned approaches can lead to pleiotropic effects, since such compounds are not highly selective. These effects are practically eliminated when using siRNA molecules. These molecules are known to be a highly accurate tool for regulation of gene expression at the post-transcriptional level. The main advantages of using siRNA molecules lie in their high efficiency at extremely low (picomolar) concentrations, minimal off-target and side-effects [10, 11]. Compared to other anti-sense strategies such as antisense DNA oligonucleotides and ribozymes, RNAi is much more potent provide an effect over a longer period of time [12].

However, the low efficiency of delivery of siRNAs to target cells due to the lack of ability of siRNA molecules to passively penetrate through the cell membrane, high probability of degradation by nucleases or through the immune system in the body hinders the implementation of the approaches under development in clinical practice and makes it necessary to search for ways to solve these problems in order to ensure effective transfection of these molecules [13, 14]. Current delivery strategies are based on the application of systems of various biological, chemical, and physical natures. Nevertheless, none of them has the properties of an ideal vector. The development of such a system over the past few years has been an urgent task of gene therapy [15]. To solve this problem, it is necessary to find methods that can ensure effective delivery of siRNAs to their target cells and do not have a significant cytotoxic effect on them [16]. The main and most well-studied signal transduction pathways leading to the induction and completion of osteogenic differentiation of MSCs are BMP/Smad and WNT/β-catenin canonical pathways. Both pathways activate the expression of the key transcription factor Runt-related transcription factor 2 (Runx2), which, in turn, initiates the expression of a cascade of marker genes associated with osteogenesis [17].

A known inhibitor of osteogenic differentiation, glycogen synthase kinase (GSK3), is a ubiquitously expressed serine/threonine kinase and has two α and β isoforms with 98% homology in their kinase domain [18, 19]. GSK inactivates glycogen synthase and was originally identified as an enzyme involved in glycogen synthesis [20]. The GSK3β isoform is most actively involved in the inhibition of the BMP and WNT/β-catenin signal transduction pathways. GSK3β is known to phosphorylate Smad1 molecules involved in the BMP pathway and β-catenin involved in the WNT/β-catenin pathway, causing their subsequent ubiquitination and proteasomal degradation [21, 22]. By this mechanism, GSK3β controls the duration of Smad1 activity as well as the translocation and accumulation of β-catenin molecules in the nucleus required for Runx2 activation. In addition, GSK3β can directly phosphorylate Runx2 and inhibits its transcriptional activity [23] (Figure 1).

**Figure 1.**
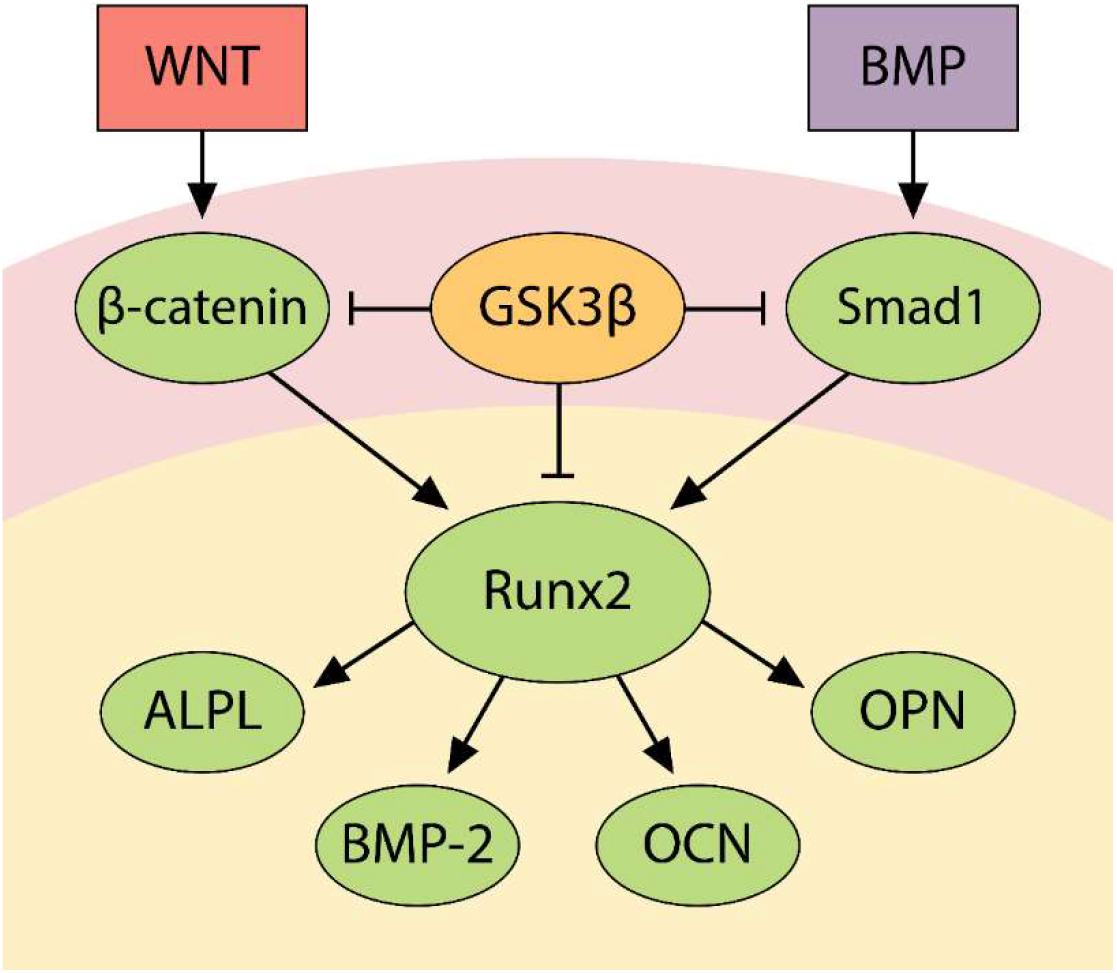
Scheme of molecular interactions between GSK3β, Smad1, and β-catenin during osteogenic differentiation.

Similarly, this kinase negatively regulates other pathways involved in the osteogenic differentiation of various mammalian cell types. GSK3β inhibits the Hedgehog signaling pathway by phosphorylating Gli1 and Gli2 proteins [18] and canonical TGF-β pathway by phosphorylating Smad3 [19] thereby preventing their translocation into the nucleus and the regulation of the transcriptional activity of *Runx2* and other osteogenic target genes [24–26].

It is known that homozygous germ line deletions of *Gsk3β* in mice causes a variable phenotype depending on genetic background, resulting in embryonic lethality or survival to day of birth with cleft palate, bifid sternum, and delayed ossification of the sternum, skull, ear bones, and cranial base. In contrast, heterozygotes display increased ossification, clavicle abnormalities, and increased bone resorption. Howewer, the *Gsk3β* insufficiency in adult mice caused an increased bone mass without other skeletal abnormalities. Authors have come to the conclusion that delayed vs increased ossification appears to depend on *Gsk3β* dosage. The phenotypes of cleidocranial dysplasia and osteoporosis in mice were significantly rescued by suppressing *Gsk3β* by oral administration of lithium chloride. Oral administration of LY603281-31-8, another low molecular weight inhibitor of this gene, increased bone formation, density and strength in an ovariectomized rat model to the levels comparable to teriparatide (human parathyroid hormone 1-34), the only osteoanabolic drug that has recently been introduced into clinical practice for osteoporosis patients. Taken together, these observations suggest that the GSK3β suppression of expression or kinase activity can provide novel therapeutics to treat bone disorders like cleidocranial dysplasia and osteoporosis (Kugimiya F. et al., 2007; Gillespie J.R. et al., 2013).

Nevertheless, there are only sporadic studies on the use of siRNA molecules to regulate osteogenic differentiation in human stem cell models [9, 27].

According to the analysis of literature data, *GSK3β* gene knockdown can shorten the terms and increase the efficiency of osteogenic differentiation of MSCs *in vitro* [7–9].

In this work we performed a comparative assessment of the efficiency of *GSK3β* mRNA knockdown and osteoinductive potential of different sequences of GSK3β siRNA molecules in the cultures of human MSCs.

To verify the efficiency of the selected siRNA molecules, we transfected and analyzed the expression of the *GSK3β* gene in HEK 293 cells. A comparative evaluation of the efficacy of lipoplexes and polyplexes formed by siRNA molecules and five commercial transfection agents most commonly used in laboratory practice on three cell lines/cultures was performed. We studied the cytotoxic effect of the most effective transfection agent in free form and as part of polyplexes on MSCs cultures.

The aim of the present study is a comparative assessment of cytotoxicity and delivery efficiency of siRNA molecules to the MSCs cultures as well as evaluation of osteoinductive properties of GSK3β siRNA molecules in these cultures.

## 2. Results

### 2.1. siRNA design

To date, there is no uniform algorithm for siRNA sequence design [35]. Freely available programs are based on different approaches (empirical rules, BLAST data, neural networks) and may produce different results when designing siRNAs even to the same sequence [36–38]. To develop our own program for designing an effective sequence of siRNA molecules, we took the empirical rules described in the review article [31] where the authors summarized a large amount of experimental data. The siRNAfit program analyzes a given sequence of 20 nucleotides, evaluating each variant of a potential siRNA for rule compliance. Conformity to a rule (e.g. “A in the third position of the sense strand”) adds a score to that sequence, while nonconformity reduces it. The top-30 siRNAs are then analyzed using BLAST software to find potential off-targets. If siRNA has a complementarity to non-target sequence for less than 16 consecutive b.p. it could be selected for synthesis [30]. As a result, two different 21-nucleotide-long siRNA sequences (conventionally designated as siRNA-3 and siRNA-4) targeting exons 7 and 8 of the GSK3β gene were designed. Also, two additional 19-nucleotides-long siRNAs (conventionally designated as siRNA-1 and siRNA-2) were selected for use described in articles [28, 29]. According to siRNAfit analysis, the siRNA-3 and siRNA-4 sequences scored the highest, 35 and 37 points, respectively. The siRNA-1 sequence scored 16 points and siRNA-2 scored 9 points.

### 2.2. Analysis of GSK3β gene knockdown efficiency in a HEK 293 cell line

To test the efficacy of GSK3β siRNA molecules, the *GSK3β* gene knockdown rate was analyzed on the model cell line HEK 293. Expression analysis of data from the FANTOM5 project [39] showed that the *GSK3β* gene is highly expressed in the HEK 293 cell line which is suitable to test efficiency of its knockdown. In addition, this cell line is characterized by high proliferative activity, which should ensure high efficiency of transfection [40]. To suppress the expression of the target gene, HEK 293 cells were transfected with GSK3β siRNA molecules 1-4 individually as well as with a mixture of all four GSK3β siRNA molecules (siRNA-multiplex). Cells of the control group were transfected with SC-siRNA molecules. The transfection agent METAFECTENE^®^ PRO and the concentration of siRNA 75 pmol/mL were chosen to increase the efficiency of siRNA delivery to HEK 293 cells. The transfection efficiency in the experiment was 84%. 24 hours after transfection with GSK3β siRNA molecules, the expression of the *GSK3β* gene was decreased by 57.05 ± 9.75% in the presence of siRNA-1, by 60.43 ± 1.44% in the presence of siRNA-2, and by 35.21 ± 13.06% in the presence of siRNA-4 relative to its expression in controls. The most effective decrease in *GSK3β* expression was caused by siRNA-3 – by 63.64 ± 11.27%. The use of an equimolar mixture of all four siRNAs was slightly less effective than the use of most siRNA molecules alone and demonstrated the decrease in expression by 56.83 ± 5.06% (Figure 2).

**Figure 2.**
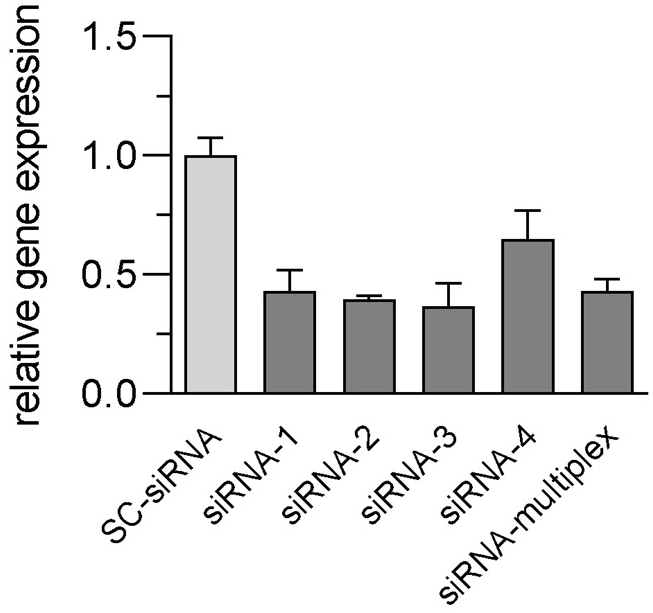
qPCR analysis of *GSK3β* gene expression in the HEK 293 cell line on day 2 after transfection with polyplexes containing 75 pmol/mL of different GSK3β siRNA molecules or a mixture thereof, or SC-siRNAs, and METAFECTENE^®^ PRO (1:2).

### 2.3. Finding the effective concentration of siRNA molecules

Using flow cytometry and fluorescence microscopy, we determined the effective concentration of siRNA molecules during transfection of AD-MSCs cells.

The METAFECTENE^®^ PRO reagent proved to be ineffective (20% or less transfected cells in culture) when preliminarily optimizing the technique of transfection of MSC cultures, while the PEI transfection agent showed the best result (data not shown). PEI was used to select the effective concentration of SC-siRNA molecules.

Polyplexes containing 50-100 pmol/mL of SC-siRNAs and PEI transfected more than 91.8% of cells in culture, whereas polyplexes containing 25 pmol/mL transfected only 55.1 ± 3.67% of cells.

It was shown that there is a direct correlation between the concentrations of siRNA, the transfection agent, and the transfection efficiency of the AD-MSCs cultures (Figure 3).

**Figure 3.**
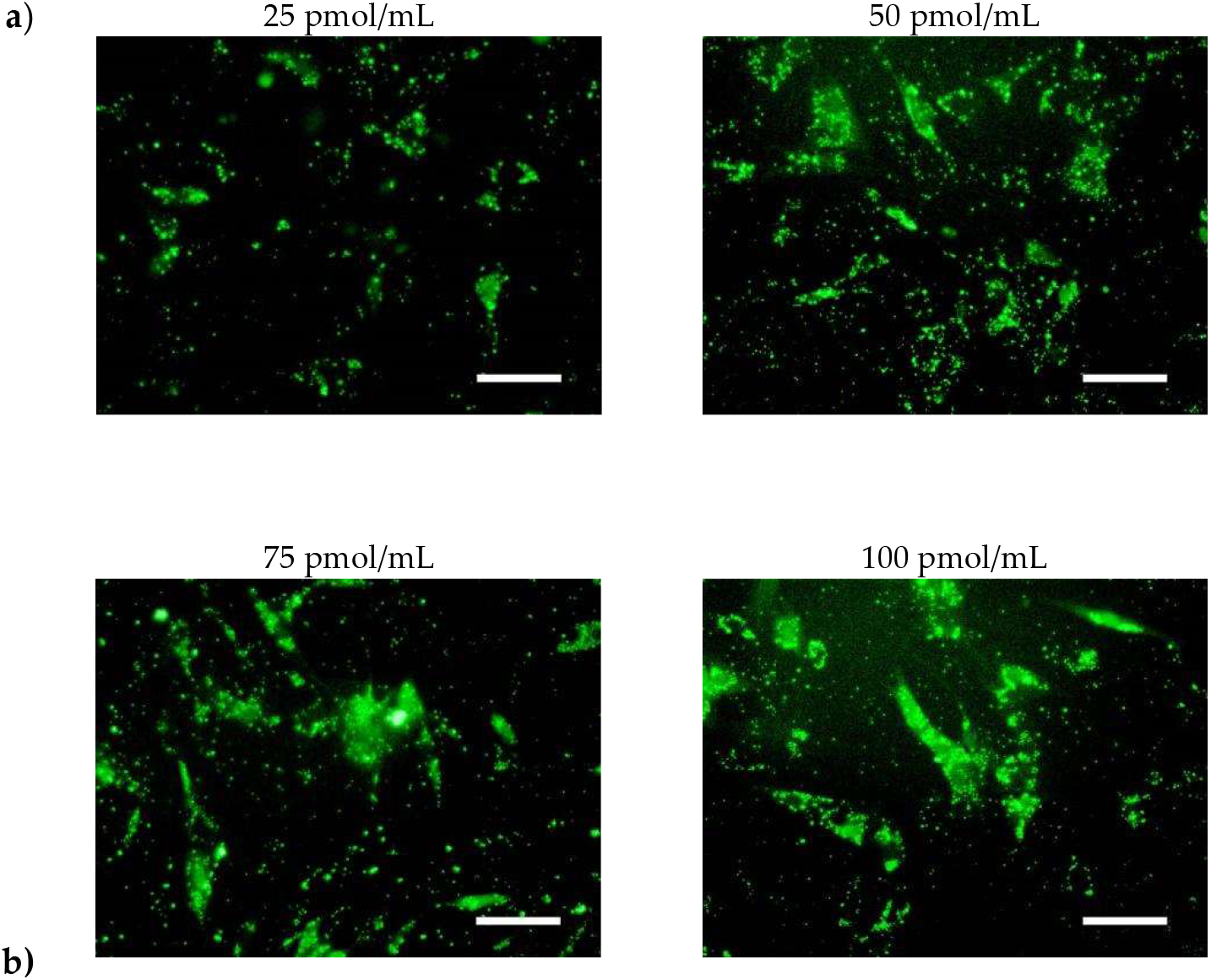
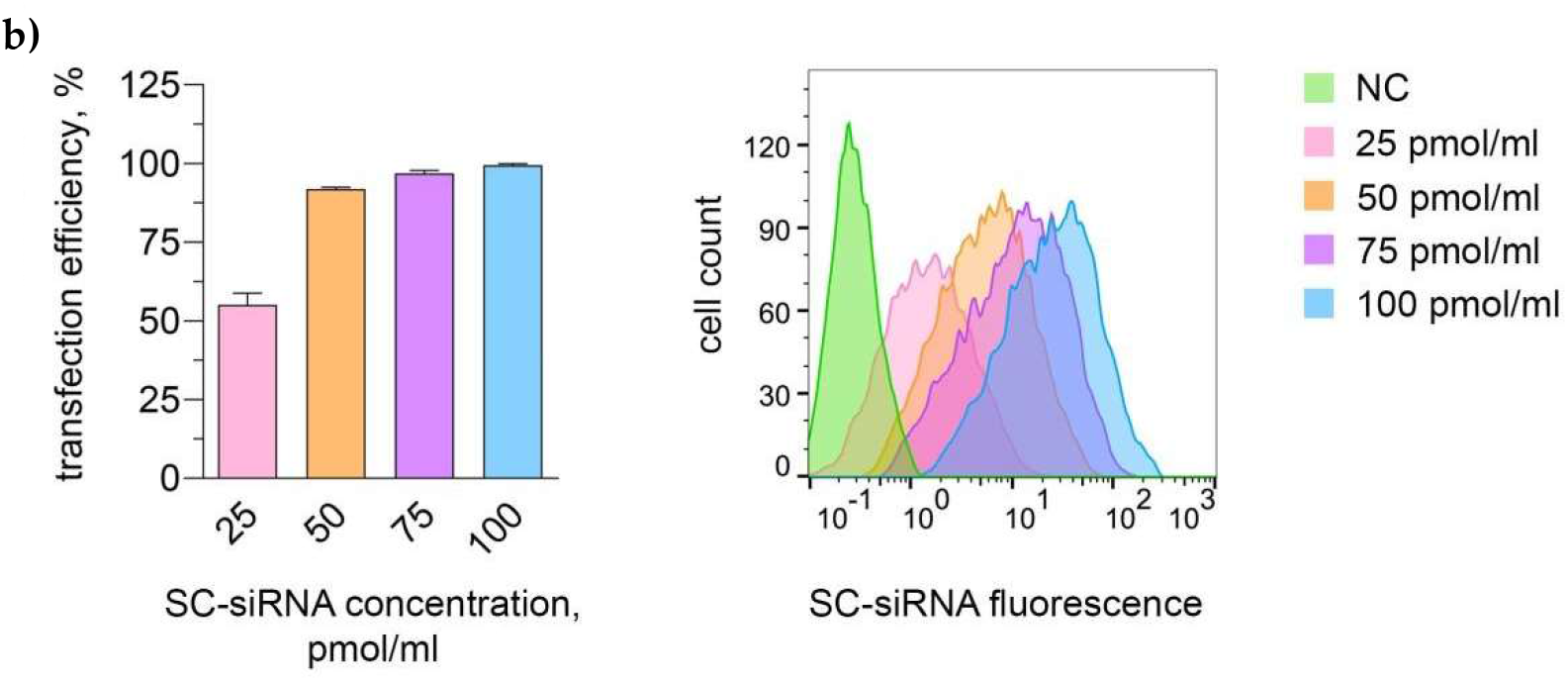
Assessment of transfection efficiency of polyplexes containing scrambled control with 6-FAM fluorescent label (SC-siRNAs) and PEI (1:3) by fluorescence microscopy (**a**) and flow cytometry (**b**). Scale bars: 100 μm. The amounts of cells analyzed are given in Appendix A.

It is known that off-target effects of siRNA molecules can be significantly reduced when they are used at relatively low doses *in vitro* [41]. Concentrations showing high transfection efficacy results (50-100 pmol/mL) in the present study are an order of magnitude lower than the concentrations causing off-target effects [41, 42].

### 2.4. Evaluation of cytotoxic effects of siRNA molecules

When SC-siRNA molecules in concentrations of 25-100 pmol/mL and GSK3β siRNA in concentrations of 50 pmol/mL were used for transfection, viability of the AD-MSCs at the end of the experiment (7 day) was at least 97.6% of the control group values. No statistically significant differences were found between the experimental and control groups, indicating the absence of cytotoxicity of SC-siRNA and siRNA GSK3β-1-4 in the selected concentrations (Figures 4, 5).

**Figure 4.**
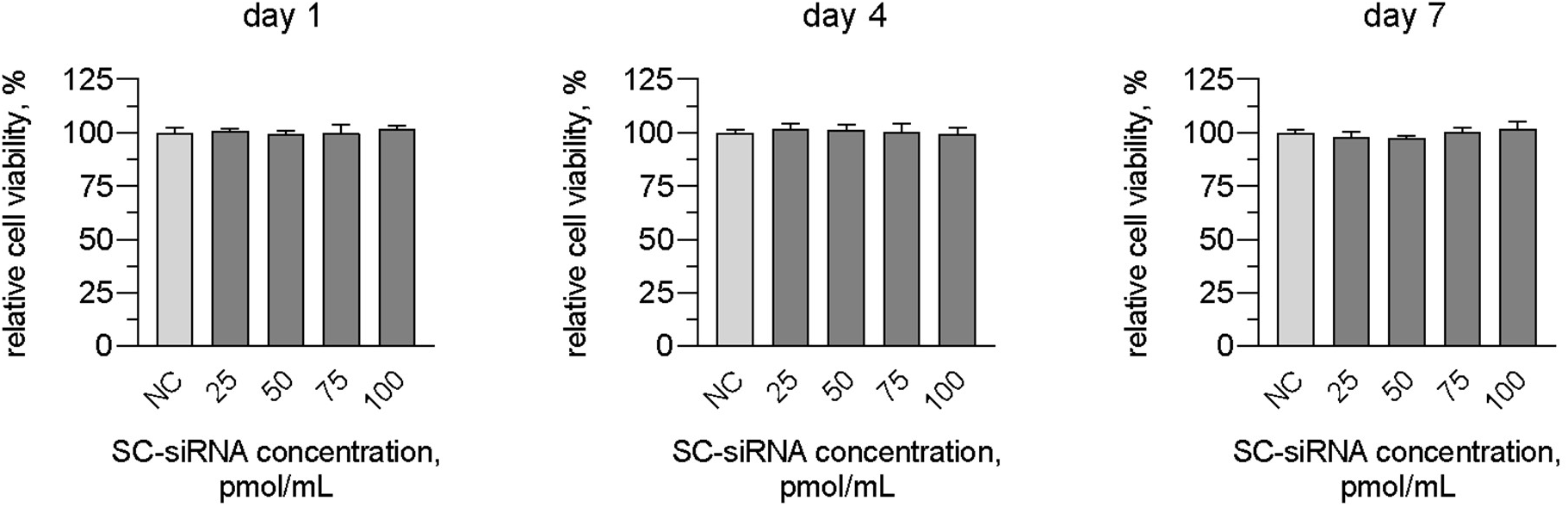
Relative viability assayed by MTT-test of AD-MSCs in the presence of 25-100 pmol/mL of SC-siRNA molecules on days 1, 4 and 7 of the experiment.

**Figure 5.**
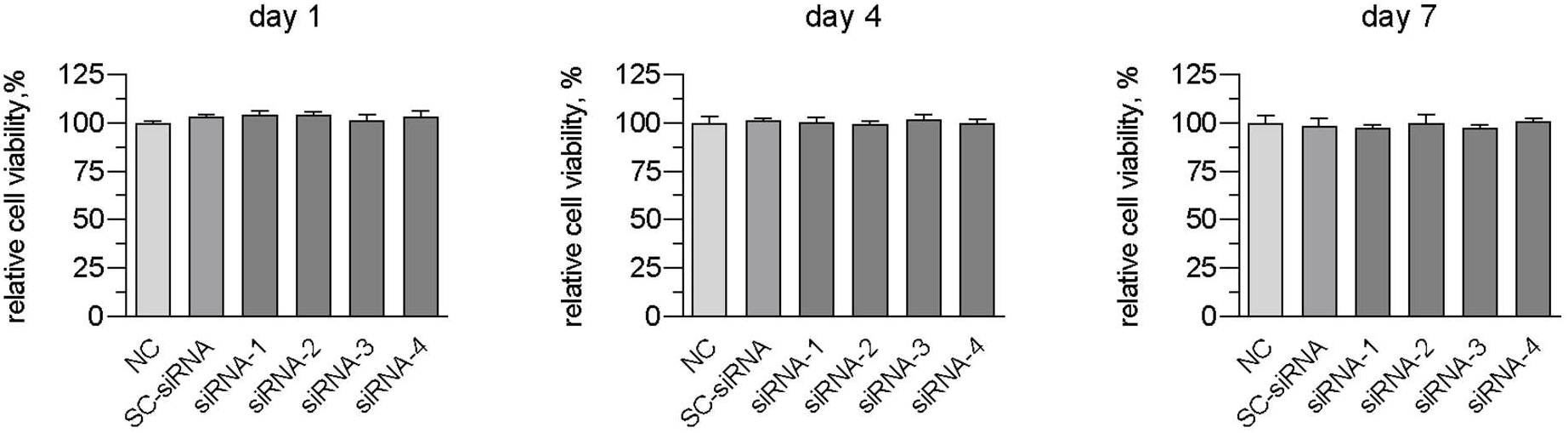
Relative viability assayed by MTT-test of AD-MSCs in the presence of 50 pmol/mL of GSK3β siRNA molecules on days 1, 4 and 7 of the experiment.

### 2.5. Evaluation of the cytotoxic effect of the PEI transfection agent

On the 1st day of the experiment the viability of the AD-MSCs in all experimental groups was more than 93.4% relative to the values of the control group. No statistically significant differences were found between the experimental and control groups.

The highest cytotoxicity was observed on days 4 and 7 of the experiment using the maximum concentrations of the transfection agent – 3 and 4 μg/mL. Cell viability relative to the control group was at least 72.3% and 75%, respectively. According to the international standard ISO 10993-5 [43], cell viability above 70% relative to the negative control indicates good biocompatibility. The use of lower concentrations (1 and 2 μg/mL) of PEI had less effect on cell survival – 81.4% and more viable cells in culture at the final term of the experiment (Figure 6).

**Figure 6.**
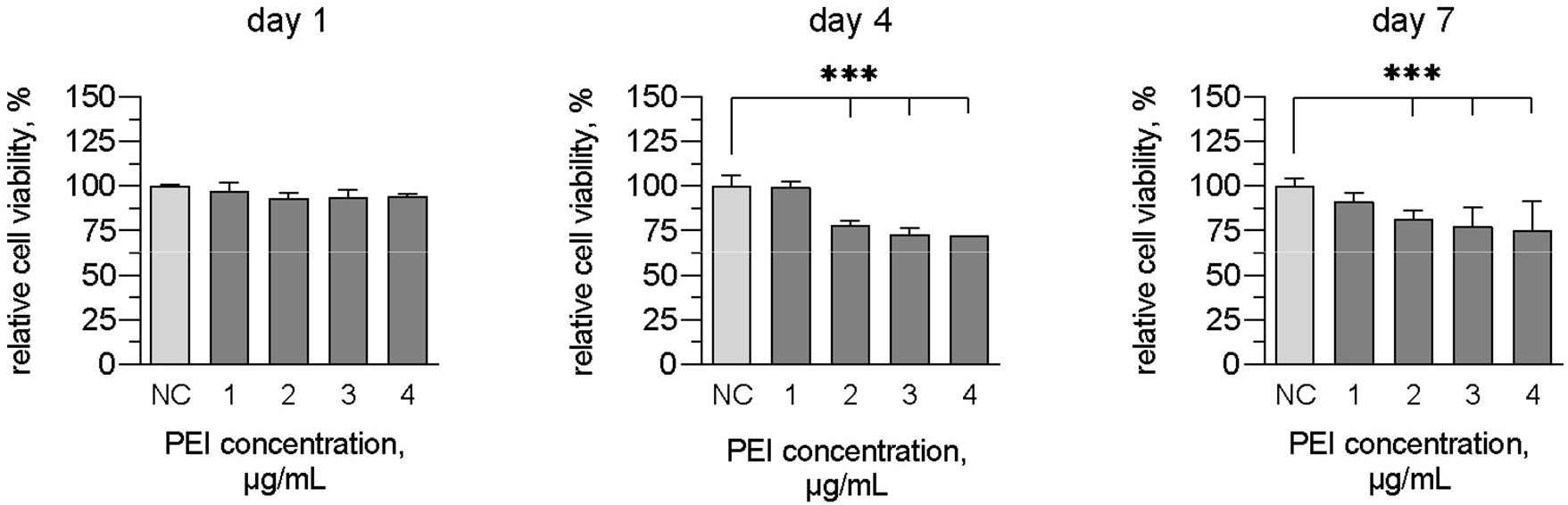
Relative viability assayed by MTT-test of AD-MSCs in the presence of 1-4 μg PEI (in a 1:3 ratio, corresponding to 25, 50, 75 and 100 pmol/mL siRNA) on days 1, 4 and 7 of the experiment.

Thus, the transfection agent PEI does not exert a marked cytotoxic effect on the AD-MSCs cultures at the concentrations of 3-4 μg/mL, corresponding to 75-100 pmol/μL of siRNA during polyplexes formation, and demonstrates biocompatibility approximating the control group at concentrations of 1-2 μg/mL, corresponding to 25-50 pmol/mL of siRNA in a 1:3 ratio (μg of siRNA : μg of PEI). An inverse relationship between the concentration of the transfection agent and cell viability is observed.

### 2.6. Evaluation of the cytotoxic effect of polyplexes containing PEI

On day 1 after transfection, all PEI-formed polyplexes, except those containing 25 pmol/mL SC-siRNAs, moderately reduced cell viability relative to the control group by no more than 20%.

On the 4th day of the experiment the viability in all experimental groups with polyplexes formed with PEI was statistically significantly different from the control (p < 0.0001) and comprised not less than 59.4% in the groups of polyplexes with the concentration of SC-siRNAs 25-75 pmol/mL and 42.3 ± 3.2% in the group with SC-siRNA concentration of 100 pmol/mL. In the presence of polyplexes containing 50 pmol/mL of SC-siRNAs and PEI, the relative viability was 70.15 ± 5.94% of cells in culture.

The most marked cytotoxic effect of polyplexes was observed on day 7 in the experimental groups with the highest SC-siRNAs and PEI concentrations. In the presence of polyplexes containing 25 pmol/mL and 50 pmol/mL of SC-siRNAs and PEI, the relative viability of cells in culture was 88.6 ± 6.48% and 60.21 ± 5.07%, respectively. Polyplexes containing 75 and 100 pmol/mL SC-siRNAs and PEI exhibited the most pronounced cytotoxicity: 33.7 ± 6.02% and 19.26 ± 2.86% of viable cells in culture (Figure 7).

**Figure 7.**
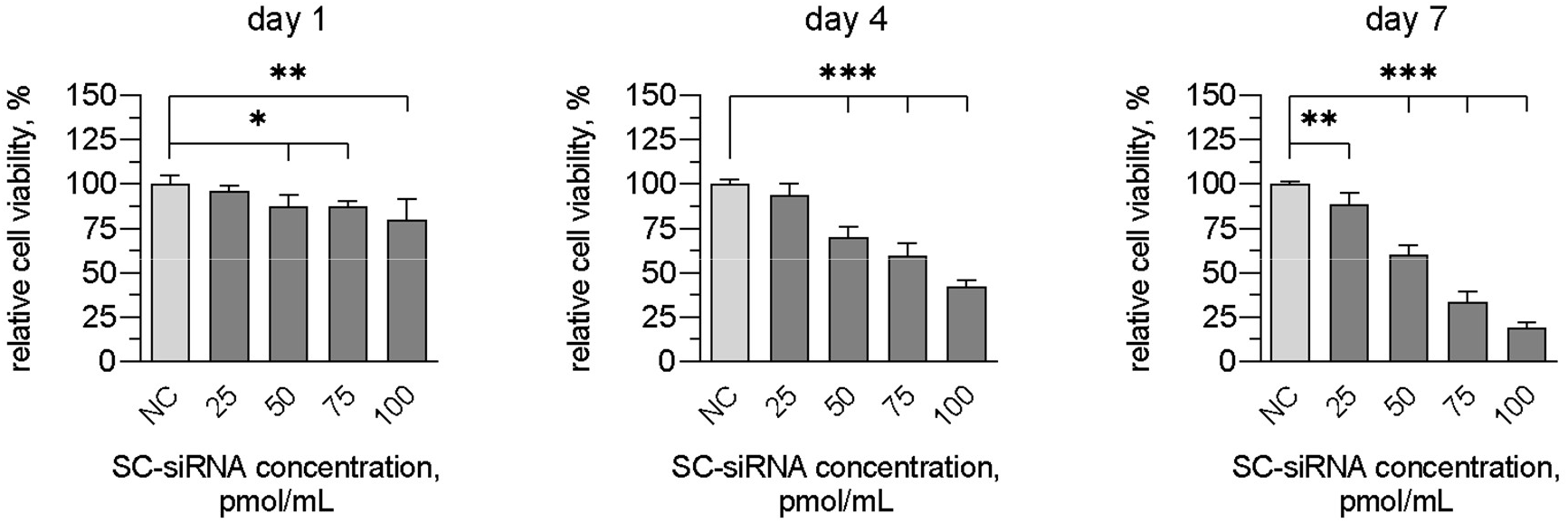
Relative viability assayed by MTT-test of the MSCs in the presence of polyplexes formed with 25-100 pmol/mL of SC-siRNA molecules and 2 μg PEI in a 1:3 ratio on days 1, 4, and 7 after transfection.

According to the ISO 10993-5 recommendations [ISO, 2009], polyplexes containing 25 pmol/mL of SC-siRNAs demonstrate high biocompatibility when incubated with MSC cultures, and polyplexes containing 50 pmol/mL – satisfactory (close to high) biocompatibility during 7 days of the experiment. Taking into account the transfection efficiency obtained using these concentrations (described in Section 2.3), the siRNA concentration of 50 pmol/mL was chosen for the following experimental works.

We can conclude that there is an inverse dependence between the concentration of siRNA and the PEI transfection agent in polyplexes and cell viability.

In addition, the transfection agent PEI and siRNA molecules separately exerted a significantly less pronounced cytotoxic effect on the AD-MSCs cultures compared to the polyplexes formed by them.

### 2.7. Choosing a transfection agent

The efficiency of transfection with lipoplexes/polyplexes formed with 50 pmol/mL of SC-siRNAs and various transfection agents on HEK 293 cell line and AD-MSCs and SHEDs cultures was studied.

The highest transfection results were achieved in HEK 293 cell line, including with liposomal agents, which confirms the available observations in the literature [40].

For the delivery of siRNA molecules into AD-MSCs and SHEDs, transfection agents based on cationic polymers were more effective than those based on liposomes. The highest transfection efficiency was shown when using the transfection agent PEI – up to 97.5% in a 1:3 ratio and up to 96.8% in a 1:2 ratio of transfected cells in MSC cultures (Figure 8). We can conclude that the efficiency of transfection with a particular reagent can vary greatly depending on the type of transfected cells (Table 1). The greatest statistical differences were observed when cells were transfected with liposome-based transfection agents. In addition, statistically significant differences were observed in the transfection of MSCs cultures obtained from different sources.

**Figure 8.**
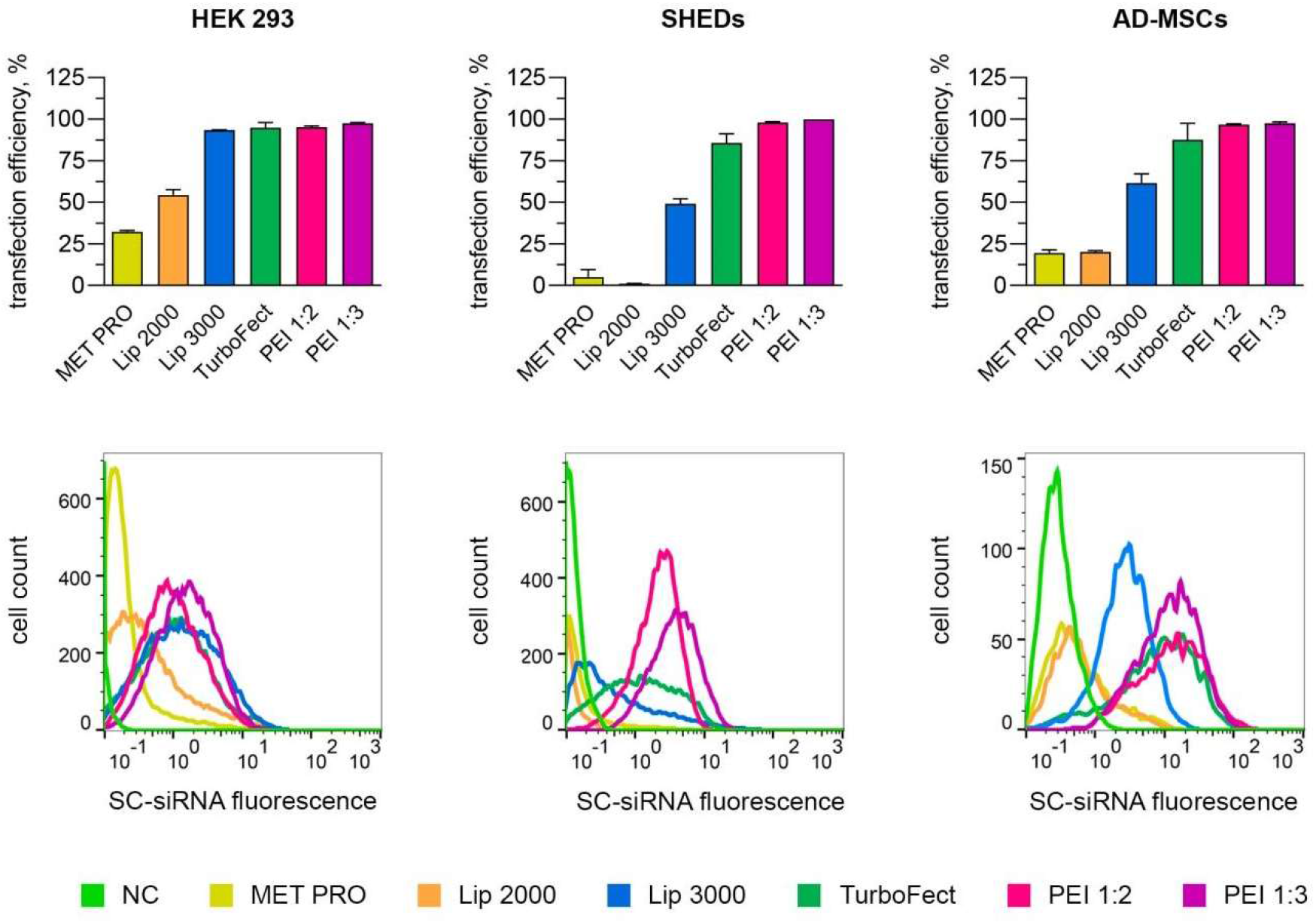
Efficiency of transfection with lipoplexes/polyplexes generated by SC-siRNAs and different transfection agents on HEK 293 cell line and cultures of human AD-MSCs and SHEDs. MET PRO: METAFECTENE^®^ PRO; Lip 2000: Lipofectamine^®^ 2000; Lip 3000: Lipofectamine^®^3000. The amounts of cells analyzed are given in Appendix B.

**Table 1.**
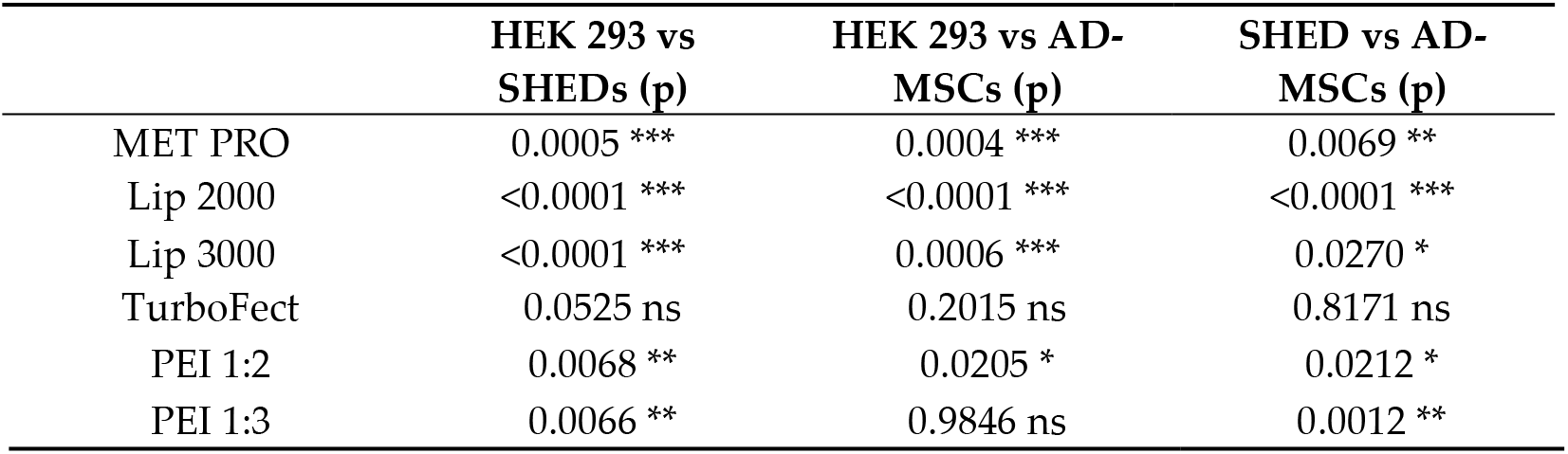
Results of a statistical analysis on an intergroup comparison of the transfection efficiency of the HEK 293 cell line and cultures of SHEDs and AD-MSCs. Unpaired Student’s t-test.

In addition, liposomal transfection agents are known to have a tendency to aggregate [44]. Liposome aggregation was observed when using METAFECTENE^®^ PRO and was not the case with Lipofectamine 2000 and Lipofectamine 3000.

### 2.8. Assessment of GSK3β gene knockdown and expression of genes of osteogenic differentiation markers in AD-MSC cultures when incubated with GSK3β siRNA

Based on the data described in the literature, to evaluate the knockdown and osteoinductive effects of GSK3β siRNA molecules in MSC cultures, the time periods 2 and 7 days after transfection were chosen [9, 45].

On the 2nd day after transfection with GSK3β siRNA molecules the *GSK3β* gene expression decreased by 20.75 ± 9.94% in the presence of siRNA-1, by 57.21 ± 1.38% in the presence of siRNA-2, and by 50.5 ± 1.96% in the presence of siRNA-4 relative to its expression in the control group. *GSK3β* expression was most effectively reduced by siRNA-3 molecules by 83 ± 1.63%.

*RUNX2* gene expression in the presence of each GSK3β siRNA increased 1.3-1.8-fold. Incubation with siRNA-3 molecules also resulted in a statistically significant increase in the expression of the other marker genes (except *OCN*): ALPL 1.5 ± 0.1-fold (p < 0.001), *BMP*-2 2.3 ± 0.2-fold (p < 0.001), *OPN* 1.2 ± 0.1-fold (p = 0.01) (Figure 9).

**Figure 9.**
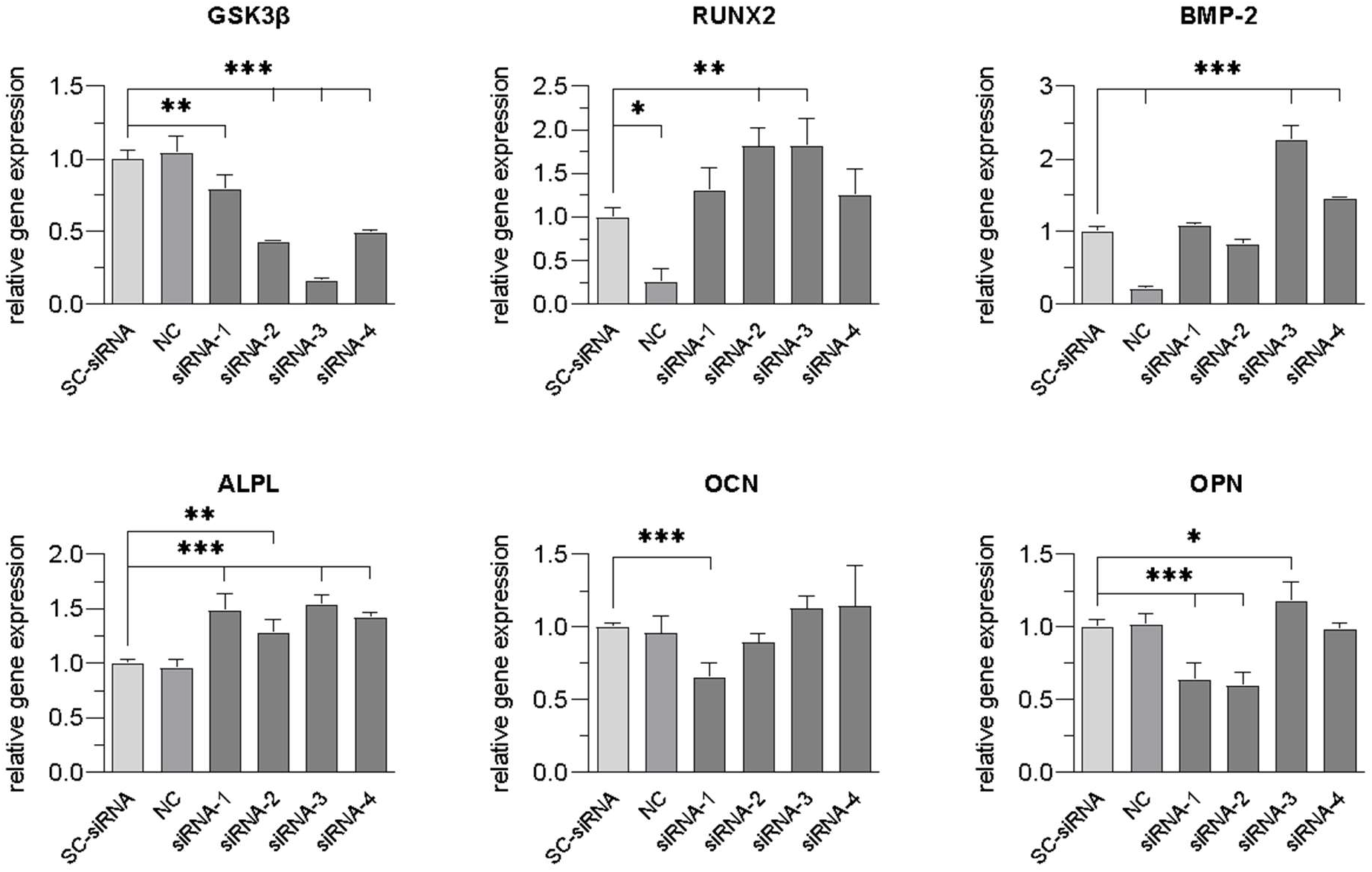
Expression of the *GSK3β* gene and genes of osteogenic differentiation marker in the AD-MSC cultures on day 2 after transfection with polyplexes containing GSK3β siRNA or SC-siRNA and PEI molecules (1:3) estimated by qRT-PCR.

On day 7 after transfection with GSK3β siRNA molecules, the expression of the *GSK3β* gene returned to normal relative to its expression in the control group. The expression of genes of osteogenic differentiation markers in the presence of siRNA-3 molecules increased 3-fold on average compared to the previous period.

The expression of *RUNX2* gene in the presence of each GSK3β siRNA molecule increased 1.9-3.8-fold in comparison with the control group, the highest increase of its expression was observed after incubation with siRNA-3 molecules – 3.8 ± 0.4 times (p < 0.001).

Incubation with siRNA-3 also resulted in the highest increase of expression of other marker genes: *ALPL* by 2.8 ± 0.4 times, *BMP*-2 by 3.7 ± 0.4 times, *OCN* by 2.3 ± 0.2 times, *OPN* by 2.7 ± 0.2 times (p < 0.001) (Figure 10).

**Figure 10.**
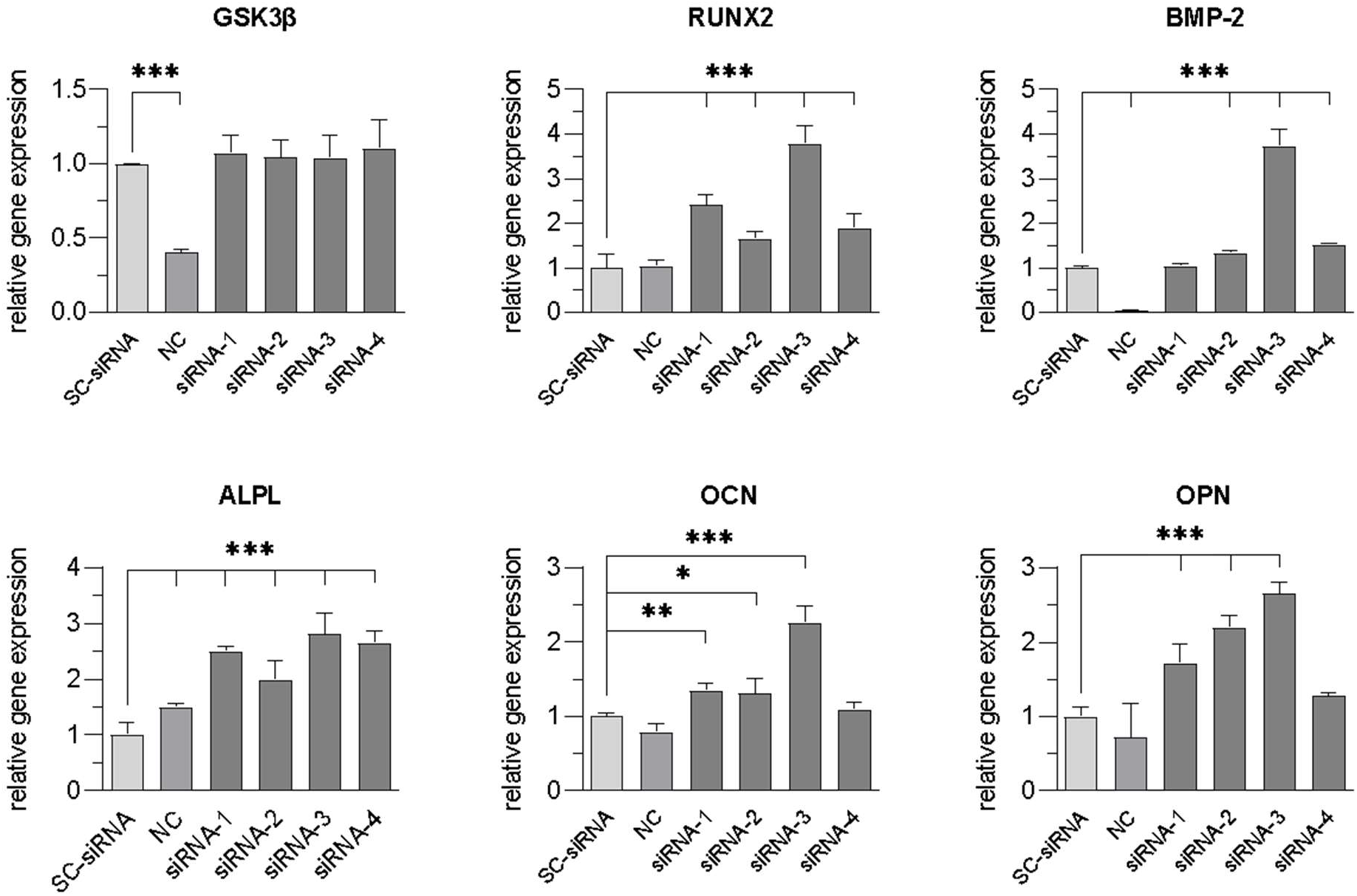
Expression of the *GSK3β* gene and genes of osteogenic differentiation markers in the AD-MSC cultures on day 7 after transfection with polyplexes containing GSK3β siRNA or SC-siRNA and PEI molecules (1:3) estimated by qPCR.

According to the results obtained, the effective knockdown of *GSK3β* gene and osteoinductive effect of GSK3β siRNA molecules at the concentration of 50 pmol/μL in the AD-MSC cultures were shown. GSK3β-3 siRNA molecules have the most marked effect.

### 2.9. Measurement of alkaline phosphatase activity and calcium ions concentrations in AD-MSC cultures after transfection with GSK3β siRNA

The transfection of AD-MSC cultures with GSK3β-3 siRNA molecules contributed to a statistically significant increase in ALPL activity and calcium ions (Ca^2+^) by day 7 of the experiment. When cell cultures were incubated with GSK3β-3 siRNA, ALPL activity and the concentration of calcium ions in cell lysates increased 6.4 ± 2.3 times (p = 0.05) and 1.3 ± 0.1 times (p < 0.0001), respectively, in comparison with the control group (Figure 11).

**Figure 11.**
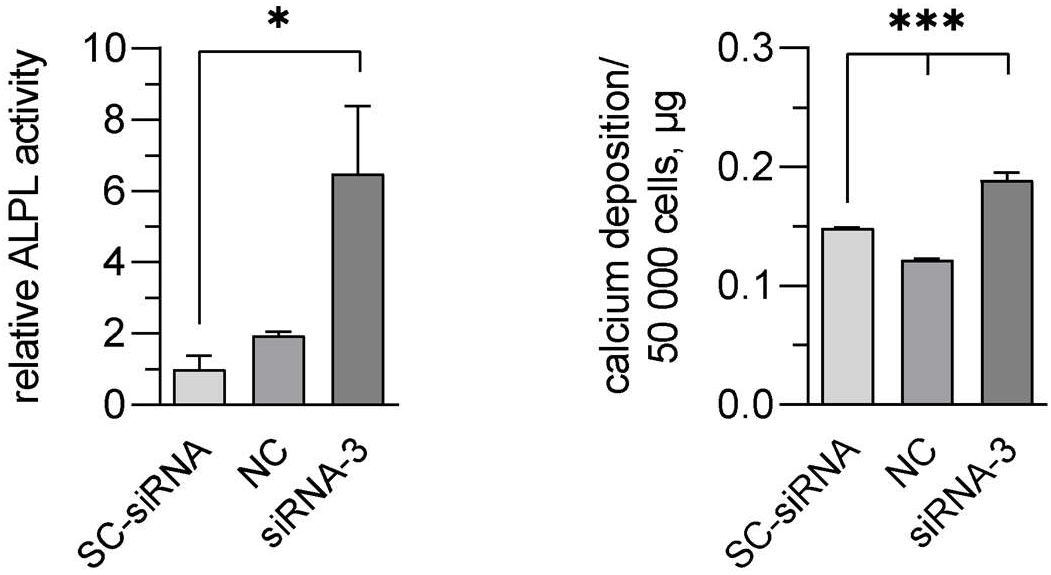
ALPL activity and Ca^2+^ concentrations in the cell lysates of AD-MSC cultures on day 7 after incubation with polyplexes containing GSK3β siRNA or SC-siRNA and PEI molecules (1:3) estimated by colorimetric assays.

Thus, the positive effect of GSK3β-3 siRNA transfection on the regulation of osteogenic differentiation of AD-MSC cultures was confirmed.

## 3. Discussion

In the works of Liang M.H., Chuang D.M., GSK3β siRNA sequences, conventionally designated as 1 and 2 in this work, or their combinations caused 70-80% reduction of the corresponding protein level 48 hours after transfection in the primary culture of rat cortical neurons. Transfection was performed using a polyamine-based chemical agent, and GSK3β knockdown was detected by Western blot [28, 29]. The results obtained in the present study at the same timeframe (reduction of GSK3β expression in HEK 293 cells using siRNA-1 and siRNA-2 by only 57 and 60%, respectively, and in MSC cultures by 21% and 57%, respectively) can be explained by the difference in the types of transfected cells, transfection agents used and methods of data detection. The sequence of siRNA-3, selected by bioinformatic methods during the present work, showed a more efficient knockdown of GSK3β gene compared to other molecules in HEK 293 and AD-MSC cultures – 64% and 83%, respectively.

According to data compiled in the siRecords database, which contains information on more than 17,000 siRNA sequences targeting Mammalian genes, less than 35% of experimentally tested siRNAs cause more than 90% gene silencing, and nearly 20% of siRNAs result in less than 50% efficiency [46, 47]. Thus, the design of the high-performance siRNA sequence is far from complete [48]. The relatively low variability of siRNA design algorithms can be explained by the limited number of siRNA sequences used during their development, as well as by the fact that these sequences were developed according to MPI principles, the Ui-Tei rule, or Reynolds’ rules [48, 49]. In this work, the authors were guided by the rules listed by Lagana A. et al., 2015 [31]. The algorithm developed uses rules that partially overlap with the Ui-Tei and Reynolds rules, but does not replicate them completely. In addition, it takes into account the presence and absence of 2-4-nucleotide motifs that empirically promote or hinder efficient RNA interference, respectively. Thus, the siRNAfit algorithm combines several approaches described by other authors and allows the design of 19-21-nucleotide siRNA sequences with a high degree of efficiency.

Chemical transfection agents may have cytotoxicity depending on the nature of their constituent components [50, 51]. This study shows that siRNA and PEI molecules separately do not exert cytotoxic effect on the MSC cultures but have moderate cytotoxicity as a part of polyplexes. Similarly, Kafil and Omidi demonstrated that polyplexes formed with linear PEI (25 kDa) and pDNA induced significantly greater cytotoxicity compared to the same transfection agent alone (free PEI) [52]. In another study, Godbey et al. showed that two types of cytotoxicity are observed when cells are transfected with PEI. These are toxicity caused by free PEI in solution and toxicity caused by PEI/pDNA polyplex processing [53]. We can conclude that the greater cytotoxicity of linear PEI as part of polyplexes compared to the transfection agent itself may be due to the processing of polyplexes inside cells.

There was also observed an inverse dependence between transfection efficiency and cell viability in the AD-MSC cultures, which is confirmed by the data of Hoare et al. In their study performed on human bone marrow-derived MSCs, increasing the amount of transfection agent based on cationic polymers led to an increase in transfection efficiency, but simultaneously significantly decreased cell viability in culture [54].

A study by Huh et al. demonstrated that inhibition of GSK3β with siRNA molecules negatively regulates osteogenic differentiation of mouse AD-MSCs [55]. However, other studies suggest that inhibition of GSK3β with siRNA molecules [9], lithium chloride [56] or other small molecules [7, 8, 57, 58] promotes osteogenic differentiation of MSCs. Wang et al. showed that GSK3β knockdown with siRNA molecules in cultures of human AD-MSCs promoted their osteogenic differentiation, leading to an increase in mRNA expression and amounts of RUNX2, Osterix, SATB2, BSP, OPN and OCN proteins, ALPL activity and amounts of calcium ions, while GSK3β overexpression significantly inhibited this process [9]. Thus, the data obtained in the present study and those of Wang et al. indicate that GSK3β acts as a negative regulator of osteogenic differentiation of human AD-MSCs, while its knockdown with the help of siRNA molecules can positively regulate the osteogenic differentiation of these cells. A novel highly efficient GSK3β siRNA sequence for the regulation of osteogenic differentiation of MSCs was developed.

## 4. Materials and Methods

### 4.1. Cell cultures

The HEK 293 line cells, human adipose tissue-derived MSCs (AD-MSCs), and MSCs derived from human exfoliated deciduous teeth (SHEDs) cultures at passages 2-3 were used in this work. MSC cultures were obtained from tissues of healthy donors who signed voluntary informed consent after obtaining the approval from the local Ethics Committee of the Research Centre for Medical Genetics (RCMG), Moscow, Russia (Protocol No.6/5 dated 15.11.2016). The ability to differentiate in adipogenic, chondrogenic and osteogenic directions, clonogenicity and immunophenotype by positive and negative cell surface CD-markers were confirmed for the MSC cultures.

Before the experiments, HEK 293 line cells and MSC cultures were cultured in DMEM medium (Paneco, Russia) containing 10% FBS (PAA Laboratories, USA), 4 mM L-glutamine (Paneco, Russia), 100 mg/L amikacin (Synthesis, Russia), 10 μg/L FGF-2 (ProSpec, USA) and 2000 IU/L heparin sodium (B. Braun Medical Inc., Germany) under standard conditions: 37 °C and 5% CO_2_. The medium was replaced every 3 days.

### 4.2. Design of siRNA molecules

In order to knockdown *GSK3β* mRNA, the sequences of 4 siRNA molecules targeting different exons of this gene were chosen. The sequences of GSK3β-1 and GSK3β-2 siRNAs were taken from the works of Liang M.H., Chuang D.M. 2006, 2007 [28, 29], the GSK3β-3 and GSK3β-4 siRNA molecules were designed using the custom-made siRNAfit program [30] developed by the Laboratory of functional genomics of the RCMG. The program scans through a given gene mRNA sequence by 19-21 b.p. windows and assigns scores to each oligonucleotide according to the empirical rules described in Lagana A., et al., 2015 [31].

Scrambled siRNA sequence labelled by 6-carboxyfluorescein (SC-siRNA) which is not targeting any human gene was taken from Horova V. et al., 2013 [32].

Sequences of all siRNAs used in the study are listed in Table 2. Synthesis of all siRNAs was performed by “DNA-Synthesis LLC” (Moscow, Russia).

**Table 2.**
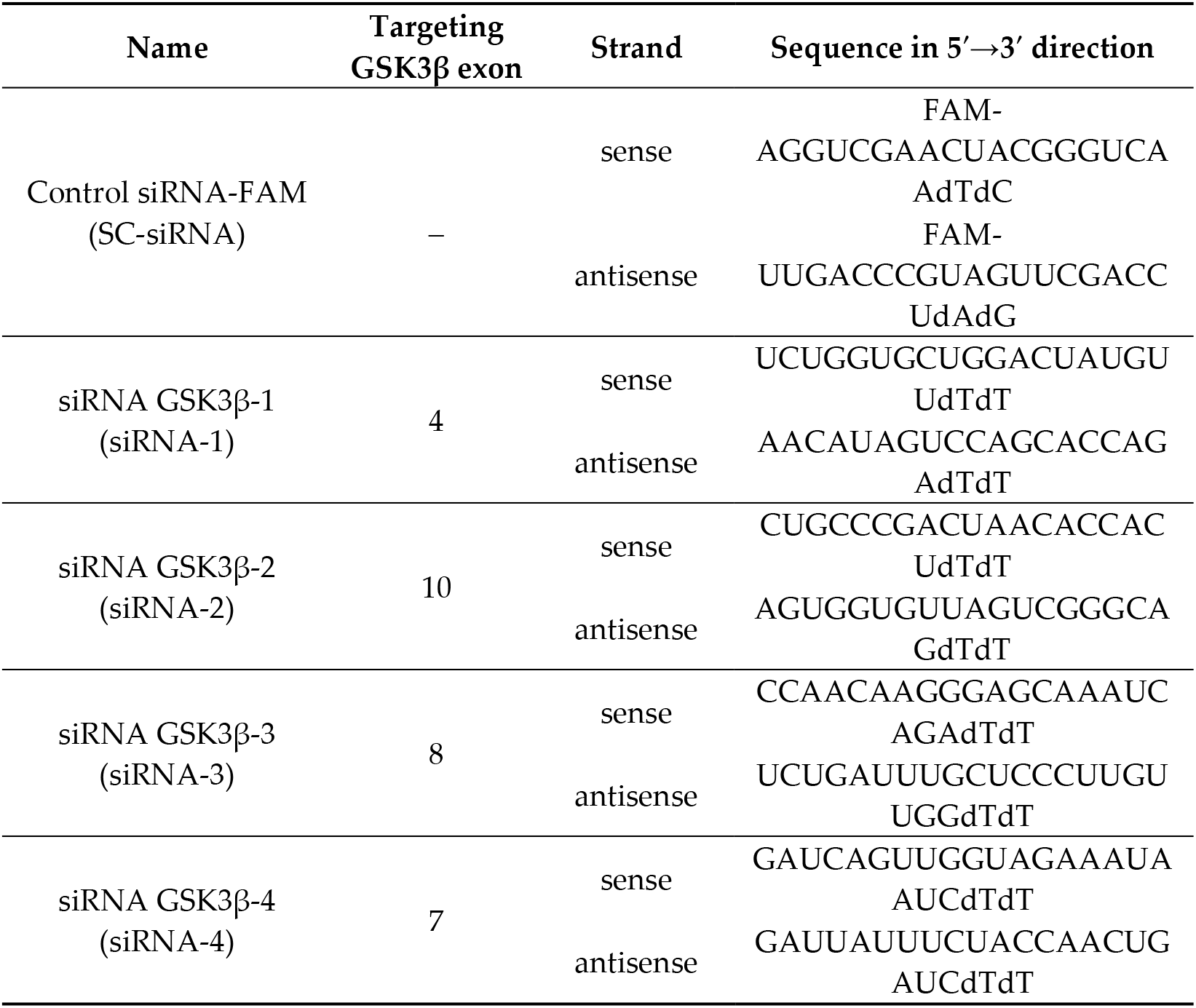
Sequences of siRNA molecules used in the study.

### 4.3. Transfection of siRNA and analysis of GSK3β gene knockdown efficiency in HEK 293 cell line

HEK 293 cell line was transfected with lipoplexes containing 75 pmol/mL of either GSK3β siRNA molecules alone or in combinations, or control SC-siRNA and METAFECTENE^®^ PRO (Biontex, USA) in a 1:2 ratio (μg of siRNA : μg or μl of transfection agent) (Table 3) according to the previously developed protocol [30]. Lipoplexes were formed in Dulbecco’s phosphate-buffered saline (DPBS) (Paneco, Russia) for 15 min. Transfection was performed in a 24-well plate when cells reached 80% confluency in 1 mL of DMEM medium supplemented with 10% FBS (PAA Laboratories, USA) for 24 h. Transfection efficiency was monitored by fluorescence microscopy and flow cytometry on the cells transfected with SC-siRNA.

**Table 3.** Quantities and ratios of siRNA and transfection agent molecules used in the experiment.

| **Volume of siRNA, $\mu\text{g}$** | **0.335** | **0.67** | **1** | **1.34** |
| --- | --- | --- | --- | --- |
| Amount of siRNA, pmol | 25 | 50 | 75 | 100 |
| Volume of DPBS/Opti-MEM for dilution of siRNA, $\mu\text{L}$ | 30 | 30 | 30 | 30 |
| Volume of DPBS/Opti-MEM for dilution of transfection agent, $\mu\text{L}$ | 40 | 40 | 40 | 40 |
| Amount of transfection agent at a ratio of 1:2, $\mu\text{g}$ or $\mu\text{L}$ | 0.67 | 1.34 | 2.01 | 2.68 |
| Amount of transfection agent at a ratio of 1:3, $\mu\text{g}$ or $\mu\text{L}$ | 1.005 | 2.01 | 3 | 4.02 |

The knockdown efficiency of the GSK3β gene in HEK 293 cells was analyzed by qRT-PCR according to the method specified in step 4.7. Total cell RNA was isolated by guanidine thiocyanate-phenol-chloroform extraction [33]. Reverse transcription was performed using the ImProm-II^™^ Reverse Transcription System reagent kit (Promega, USA) according to the manufacturer’s recommendations. The primer sequences for *GSK3β* mRNA are given in Section 4.7 and Table 4. Primer sequences for reference genes are given in the article [30].

**Table 4.**
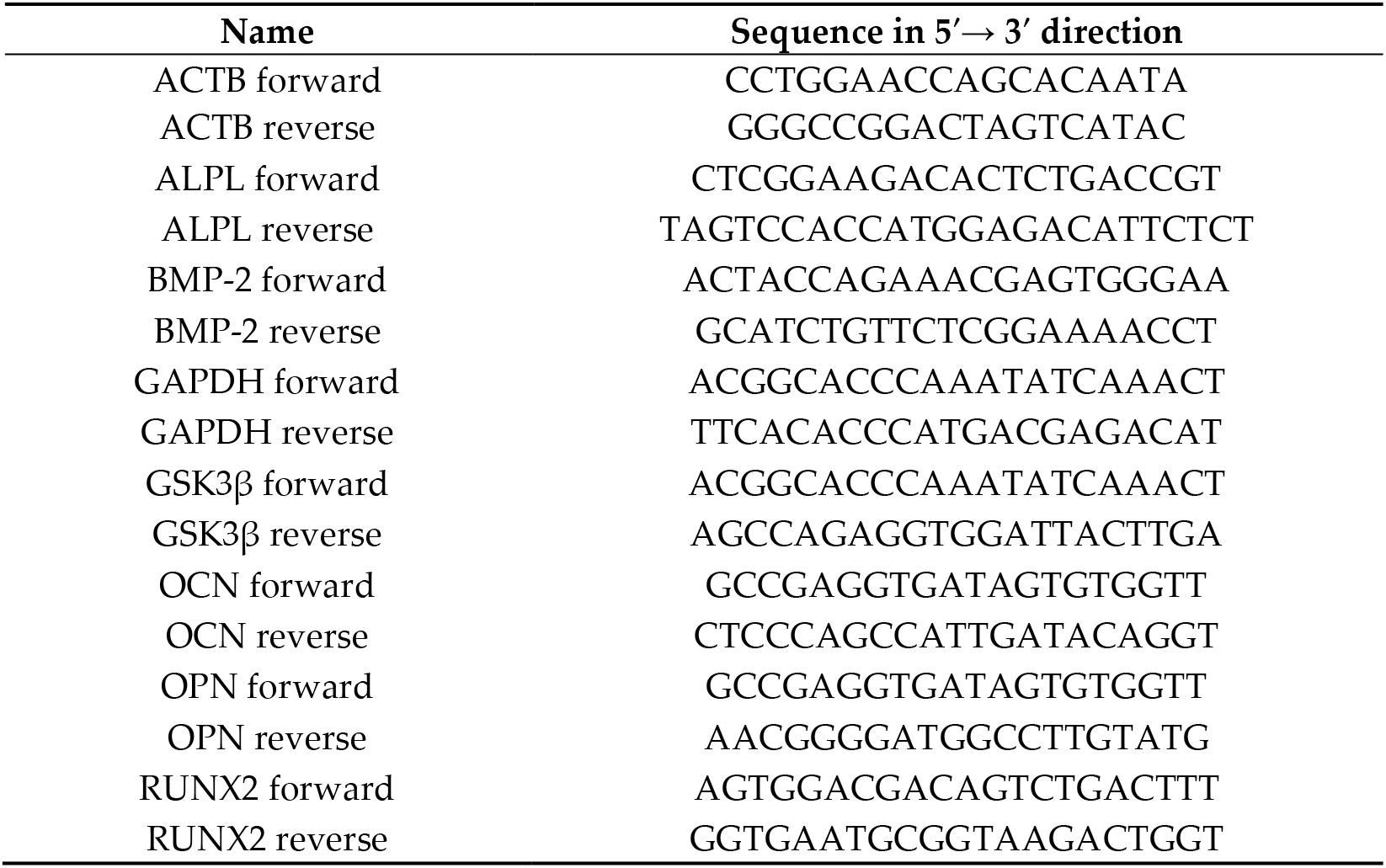
Sequences of primers specific to the *GSK3β* gene, genes of osteogenic differentiation marker, and reference genes.

The experiments by fluorescence microscopy were performed in 2 biological and 5 technical replicates. qPCR-RT analysis was performed in 3 technical replicates for each group.

### 4.4. Transfection of siRNA molecules into MSC cultures

Interventionary studies involving animals or humans, and other studies that require ethical approval, must list the authority that provided approval and the corresponding ethical approval code.

To determine the effective concentration of siRNA molecules AD-MSCs were transfected with polyplexes containing 25, 50, 75, or 100 pmol/mL of SC-siRNA and linear polyethylenimine (PEI) with a molecular weight of 25 kDa (23966-1; Polyscience, USA) in a 1:3 ratio (μg of siRNA : μg or μL of transfection agent) (Table 3).

In order to choose the optimal transfection agent, the HEK 293 cell line and MSC cultures were transfected with lipoplexes/polyplexes consisting of 50 pmol/μL of SC-siRNA and METAFECTENE^®^ PRO (Biontex, USA), Lipofectamine^®^ 2000, Lipofectamine^®^3000 with addition of “P3000”, TurboFect (Thermo Fisher Scientific, USA) reagent in the ratio 1:2; or PEI at a ratio of 1:2 or 1:3 according to the manufacturers’ recommendations (Table 3).

Lipoplexes/polyplexes were formed in DPBS or Opti-MEM medium (Thermo Fisher Scientific, USA) for 35-40 min. Transfection was performed in a 24-well culture plate when cells reached 70-80% confluency in 1 mL Opti-MEM medium with 5% FBS (PAA Laboratories, USA) for 24 hours.

Cells that were incubated in equivalent amounts of Opti-MEM medium with 5% FBS, and addition of DPBS or Opti-MEM were used as controls.

Digital images of transfected cells at the bottom of culture plates were obtained and processed using an inverted microscope AG Axio Observer D1 with the possibility of fluorescence detection and AxioCam HRc camera, as well as ZEN software (Carl Zeiss Microscopy GmbH, Germany). The efficiency of cell transfection in suspension was assessed using a CyFlow^®^ Space Flow Cytometer (Partec, USA) and FloMax^®^Software package (Partec, USA). Combined charts were plotted using FlowJo^™^ Software v10.7.1 (BD Biosciences-Europe, USA).

The experiment was conducted in 5-8 biological and 1 technical replicates for each group.

### 4.5. Selection of cytocompatible concentration of siRNA molecules and transfection agent in polyplexes

To select the optimal concentration of siRNA molecules and transfection agent in polyplexes, we performed a comparative assessment of their cytotoxic effect on AD-MSC cultures for 7 days after 24 hours of incubation in 24-well plates in 1 mL of Opti-MEM medium supplemented with 5% FBS.

To evaluate the cytotoxic effect of siRNA molecules, cells were incubated with SC-siRNA or siRNA GSK3β at concentrations of 25, 50, 75, or 100 pmol/μL.

To evaluate the cytotoxic effect of transfection agents, cultures were incubated with 0.67 to 2.68 μL/mL TurboFect or 1 to 4 μg/mL PEI (Table 3), which corresponds to a ratio of 1:2 (μg siRNA : μL TurboFect) and 1:3 (μg siRNA : μg PEI) for formation of polyplexes with concentrations of 25, 50, 75, or 100 pmol/μL of siRNA molecules.

The cytotoxicity of polyplexes containing 25, 50, 75, or 100 pmol/μL of SC-siRNA and TurboFect or PEI at a ratio of 1:2 and 1:3, respectively, was then examined (Table 3).

Immediately after transfection and every third day, the cultural media were replaced with DMEM medium (Paneco, Russia) supplemented with 10% FBS, 4 mM L-glutamine (Paneco), and 100 mg/L amikacin (Sintez, Russia). Cytotoxicity of siRNA molecules, transfection agents and polyplexes were analyzed on days 1, 4-5 and 7 after transfection using MTT-test according to the standard technique [34] which determines the number of living cells in a well of a plate.

#### 4.5.1. MTT-test

MTT (Paneco, Russia) was added to the cell cultures at a concentration of 0.5 mg/mL and incubated for 2.5 h at 37 C. Formazan crystals were extracted from the cells with 500 μL dimethyl sulfoxide (PanEco, Russia) by stirring the suspension on an orbital thermoshaker for 20 min. Formazan absorbance was measured on an EnSpire^®^ Multimode Plate Reader (PerkinElmer, USA) at 570 nm and the background value at 620 nm was subtracted.

The experiment was conducted in 4-5 biological and 1 technical replicates for each group.

### 4.6. Assessment of GSK3β gene knockdown efficiency and osteoinductive properties of GSK3β siRNA molecules

For *GSK3β* gene knockdown and osteogenic differentiation, transfection of AD-MSCs with polyplexes formed with 50 pmol/mL of GSK3β siRNA and PEI in a 1:3 ratio (μg siRNA : μg PEI) according to the procedure described in Section 2.3 was performed.

As a negative control, AD-MSCs transfected with SC-siRNA and AD-MSCs cultured in equivalent to the experimental groups amounts of Opti-MEM medium supplemented with 5% FBS or addition of DPBS.

Immediately after transfection and every third day, the cultures media were replaced with DMEM (Paneco, Russia) supplemented with 10% FBS, 4 mM L-glutamine (Paneco), and 100 mg/L amikacin (Synthesis, Russia).

Transfection results were considered positive when the efficiency of transfection with SC-siRNA molecules exceeds 90%. In that case, the expression of *GSK3β* gene and genes of osteogenic differentiation markers was evaluated on days 2 and 7 of the experiment, the activity of alkaline phosphatase and calcium ions amount were also determined.

The experiments by fluorescence microscopy were performed in 3 biological and 3 technical replicates. qPCR-RT analysis was performed in 3 technical replicates for each group.

### 4.7. Evaluation of expression of GSK3β gene and genes of osteogenic differentiation marker

Total RNA was isolated from MSCs cultures using RNeasy Plus Mini Kit (Quagen, Germany), the synthesis of the first chain of total cDNA on RNA matrix was performed using RevertAid kit (Thermo Scientific, Germany) according to the manufacturers’recommendations.

qRT-PCR was performed using a CFX96 Touch^™^ thermal cycler (Bio-Rad, USA) using intercalating SYBR Green I dye (Eurogen, Russia) and primers specific to the *GSK3β* gene and genes of osteogenic differentiation marker: *ALPL*, *RUNX2*, *BMP-2*, *OCN*, *OPN*. The mRNA expression level of the analyzed genes was normalized to the average expression values of the *GAPDH* and *ACTB* reference genes (Table 4).

The protocol of qPCR included the following steps: denaturation at 95 °C for 6 min followed by 45 cycles of denaturation at 95 °C for 10 s, primer annealing at 60 °C for 15 s, elongation at 72 °C for 20 s. The specificity of qPCR was confirmed by melting curve analysis (from 70 to 98 °C, with 0.5 °C increments in each cycle) and by PAGE. The obtained expression results of the analyzed genes were normalized to the average expression values of the reference genes *GAPDH* and *ACTB* in the same sample. The relative expression of the gene of interest was calculated by the 2^-ΔΔCt^ method.

### 4.8. Estimation of alkaline phosphatase activity and calcium ions concentration

Activity of ALPL and calcium ions concentration in cell lysates on day 7 of osteogenic differentiation were determined using reagent kits “Alkaline Phosphatase-New Liquid Form” (Vector-Best, Russia) and Calcium Colorimetric Assay Kit (Sigma-Aldrich, USA), respectively, according to the manufacturers’ instructions, on an EnSpire Multimode Plate Reader (PerkinElmer, USA).

### 4.9. Statistical analysis

Statistical processing of the data and graphing was performed using GraphPad Prism 8.00 software (USA). Intergroup differences were determined using Holm-Šidák test when comparing 3 or more groups and unpaired Student’s t-test when comparing 2 groups. Differences with p-values ≤ 0.05 were considered statistically significant. Significance of statistical differences for the experimental groups relative to controls at *p* ≤ 0.05 is designated as “*”, *p* < 0.01 – as “**”and *p* < 0.001 – as “***” throughout the manuscript.

## 5. Conclusions

siRNA molecules, acting by the RNA interference mechanism, are a highly accurate tool for genetic silencing of target mRNA transcripts without directly acting on the genome. However, the chemical transfection agents used for their delivery can have cytotoxicity [50, 51]. In the course of this work, we determined the most effective agent for transfection of MSCs – PEI and the optimal concentration of siRNA molecules in polyplexes – 50 pmol/mL. This transfection agent allows transfection of up to 97.5% of cells in MSC cultures. siRNA molecules and transfection agents separately do not have cytotoxic effect on MSC cultures, but have moderate cytotoxicity in polyplexes. Moreover, there is a direct correlation between the concentrations of siRNA, the transfection agent, and the transfection efficiency of these cultures. Also, there is an inverse relationship between the concentration of siRNA and the transfection agent in the polyplexes and cell viability. Finally, the pronounced osteoinductive properties of GSK3β siRNA molecules in human MSC cultures were demonstrated. The developed GSK3β-3 siRNA sequence showed the best result of target gene knockdown and osteoinductive effect simultaneously. Due to the ability to positively regulate the osteogenic differentiation of MSCs, this sequence can be recommended for research purposes for the induction and enhancement of the osteogenic differentiation of MSCs.

In addition, the results obtained can be applied in the development of gene therapy strategies based on siRNA molecules in human bone tissue diseases.

## Author Contributions

Conceptualization, Elena Galitsyna, Tatiana Bukharova, Mikhail Skoblov and Dmitriy Goldshtein; Data curation, Irina Krivosheeva and Mikhail Skoblov; Formal analysis, Elena Galitsyna, Anastasiia Buianova, Irina Krivosheeva and Mikhail Skoblov; Funding acquisition, Mikhail Skoblov and Dmitriy Goldshtein; Investigation, Elena Galitsyna, Anastasiia Buianova and Irina Krivosheeva; Methodology, Elena Galitsyna, Irina Krivosheeva and Mikhail Skoblov; Project administration, Elena Galitsyna, Tatiana Bukharova, Mikhail Skoblov and Dmitriy Goldshtein; Software, Irina Krivosheeva and Mikhail Skoblov; Supervision, Elena Galitsyna, Tatiana Bukharova, Mikhail Skoblov and Dmitriy Goldshtein; Validation, Elena Galitsyna, Anastasiia Buianova and Irina Krivosheeva; Visualization, Elena Galitsyna and Anastasiia Buianova; Writing – original draft, Elena Galitsyna; Writing – review & editing, Tatiana Bukharova and Mikhail Skoblov. All authors have read and agreed to the published version of the manuscript.

## Institutional Review Board Statement

The study was conducted according to the guidelines of the Declaration of Helsinki, and approved by the local Ethics Committee) of the Research Centre for Medical Genetics (RCMG), Moscow, Russia (Protocol No.6/5 dated 15.11.2016).

## Informed Consent Statement

Informed consent was obtained from all subjects involved in the study.

## Acknowledgments

The authors are grateful for the help with the translation of this article into English to Dr. Andrey Marakhonov, a Senior Researcher of the Laboratory of functional genomics of the RCMG.

## Abbreviations

6-FAM: 6-carboxyfluorescein
ACTB: actin beta
AD-MSCs: human adipose tissue-derived mesenchymal stem cells
ALPL: alkaline phosphatase
BMP-2: human bone morphogenetic protein-2
DMEM: Dulbecco’s modified Eagle’s medium
DPBS: Dulbecco’s phosphate-buffered saline
FBS: fetal bovine serum
GAPDH: glyceraldehyde 3-phosphate dehydrogenase
GSK3β: glycogen synthase kinase-3β
MSCs: mesenchymal stem cells
OCN: osteocalcin
OPN: osteopontin
PAGE: polyacrylamide gel electrophoresis
pDNA: plasmid DNA
PEI: polyethylenimine
qPCR: quantitative real-time polymerase chain reaction
Runx2: runt-related transcription factor 2
SC-siRNA: scrambled siRNA sequence labelled by 6-carboxyfluorescein
SHEDs: mesenchymal stem cells derived from human exfoliated deciduous teeth
siRNA: small interfering RNA.

## Appendix A

**Table A1.**
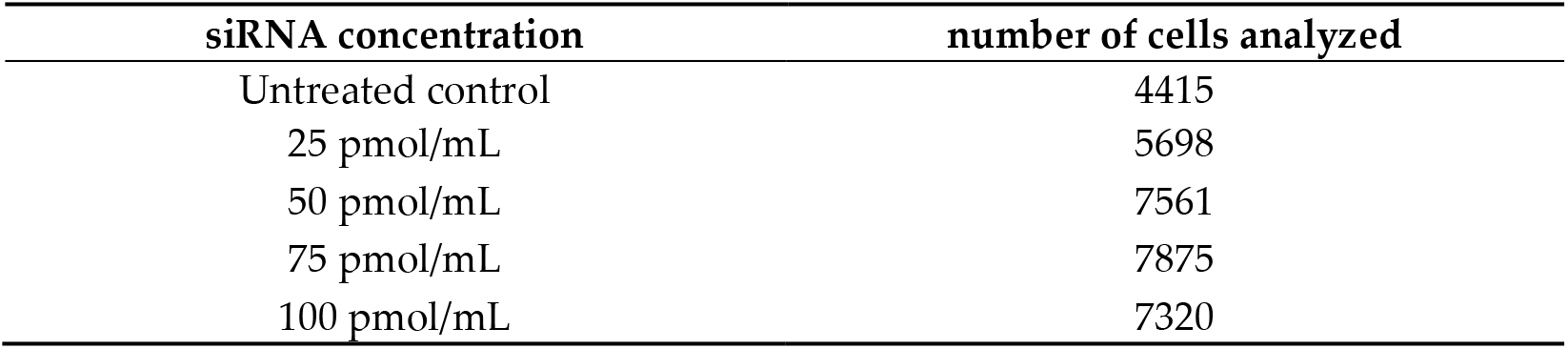
The number of cells analyzed when evaluating the efficiency of transfection with polyplexes containing control SC-siRNA and PEI (1:3) by flow cytometry.

**Table A2.**
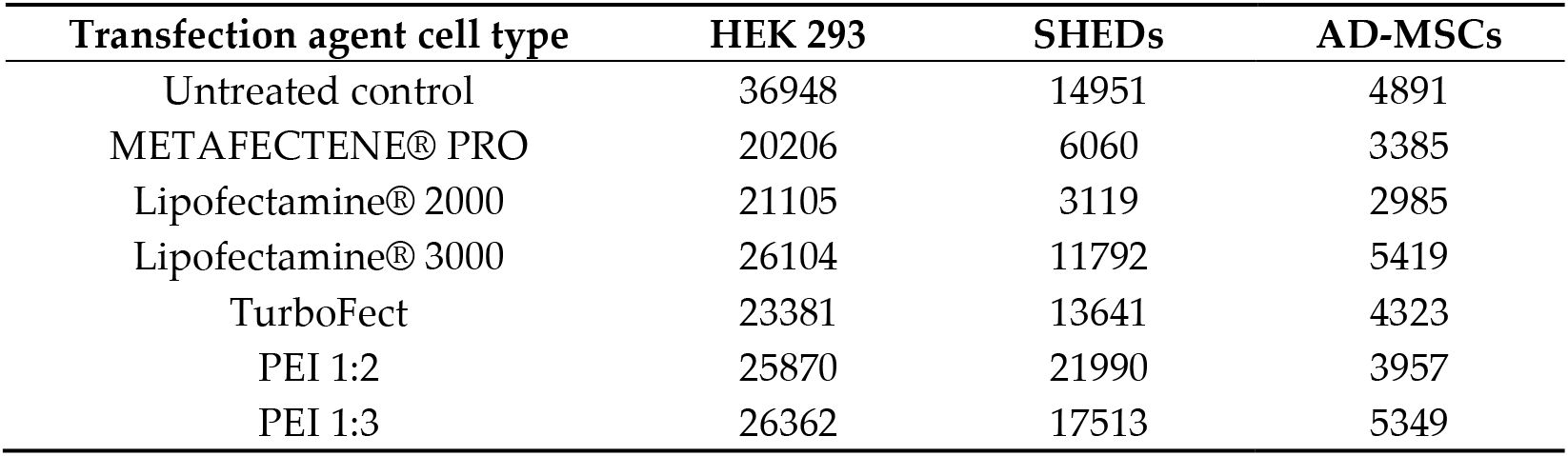
The number of cells analyzed in the evaluation of transfection efficiency of lipoplexes/polyplexes formed with different transfection agents on HEK 293 cells and human AD-MSCs and SHEDs cultures.

